# Prevalent uses and evolution of exonic regulatory sequences in the human genome

**DOI:** 10.1101/2021.09.06.459186

**Authors:** Jing Chen, Pengyu Ni, Meng Niu, Jun-tao Guo, Zhengsheng Su

## Abstract

**Background:** It has long been known that exons can be used as *cis-*regulatory sequences such as enhancers. However, the prevalence of such dual-use of exons and how they evolve remain elusive. Our recently predicted highly accurate, large sets of *cis*-regulatory module candidates (CRMCs) and non-CRMCs in the human genome positioned us to address these questions.

**Results:** We found that exonic transcription factor binding sites (TFBSs) occupied at least a third of the total exon lengths, and 96.7% of genes had exonic TFBSs. Both A/T and C/G in exonic TFBSs are more likely under evolutionary constraints than those in non-CRMC exons. Interestingly, exonic TFBSs in codons tend to encode loops rather than more critical helices and strands in protein structures, while exonic TFBSs in untranslated regions (UTRs) tend to avoid positions where known UTR-related functions are located. Moreover, active exonic TFBSs tend to be in close physical proximity to distal promoters whose immediately downstream genes have elevated transcription levels, suggesting that they might be involved in transcriptional regulation of target genes. It is highly possible that less critical positions in an exon that is physically close to a promoter can evolve into a TFBS when no non-exonic sequences are physically available to the promoter.

**Conclusions:** Exonic TFBSs might be more prevalent than originally thought and are likely in dual-use. Possible detrimental effects caused by such dual-use can be reduced by using less critical exonic positions. We proposed a parsimonious model to explain how a stretch of codons evolve into a TFBS.

## Background

Eukaryotic genomes contain two major types of functional sequences: genes that encode proteins or RNAs, and *cis*-regulatory sequences that control the transcription of the genes [1–3]. *Cis*-regulatory sequences including promoters, enhancers, silencers and insulators are also known as *cis*-regulatory modules. A CRM is typically made up of closely located transcription factor (TF) binding sites (TFBSs) of the same and different TFs along a segment of DNA with varying lengths ranging from a few hundred to a few thousand base pairs [4]. The binding of cognate TFs to the TFBSs in a CRM may lead to changes in the transcriptional rate of the target gene in a cell/tissue type specific manner [4]. CRMs have long been known to reside in intergenic regions or in the introns of genes [5–8]. However, it has been found that exons including protein coding DNA sequences (CDSs), 5’-untranslated regions (5’-UTRs) and 3’-UTRs, could also function as distal CRMs, such as enhancers and silencers for their own or different genes [9–25]. For example, it has been shown that the second exon of the human ADAMTS5 gene regulates its own transcription [11], and the 15^th^ exon of mouse gene Dync1i1 physically interacts with and activates the promoter of the *Dlx5/6* gene 900kb away [13]. Mutations in such exonic enhancers can result in diseases [15, 26–29]. More recently, it has been reported that at least 15% of codons in 86% of genes in the human genome were hypersensitive to DNase I treatment in 81 human cell/tissue types and thus were likely in dual-use for encoding amino acid sequences in proteins and TFBSs in DNA for gene transcriptional regulation [30]. These so-called duons were found to be more conserved than non-duon codons at the four-fold degenerate codon positions, and were thought to be under double negative selections owing to their dual-use [30]. Cases have been found that mutations in duons could lead to diseases due to altered activities of relevant exonic enhancers [31].

Nonetheless, a reanalysis of the DNase I hypersensitive sites (DHSs) showed that duons evolved similarly to non-duon codons when the conservation levels or substitution rates of A/T and C/G were compared separately, using either phyloP [32] scores or substitution rates derived from the most recent common ancestor of humans and gorilla compared with chimpanzee. The authors [30] argued that the earlier conclusions about duons were incorrectly drawn due to the higher C/G frequencies at the fourfold degenerate codon positions in duons than in non-duon CDSs, whereas C/G at the fourfold degenerate codon positions tend to be more conserved than A/T. This result casts doubts on the validity of the duon hypothesis [33]. A similar conclusion was drawn based on an analysis of three known exonic enhancers [34]. On the other hand, we [35] and others [36, 37] have shown that DHSs are not a reliable predictor of CRMs, not to mention more subtle TFBSs in CRMs, because a naked DNA segment hypersensitive to DNase I treatment may not necessarily be a CRM [36–39]. It is highly likely that the earlier predicted duons might have a high false discovery rate (FDRs) [35–39]. Therefore, the evolutionary neutrality of the duons observed earlier [33] might be due to inaccuracy of duons predicted based on DHSs. To clarify these contradictory results, ideally, one might need a sufficiently large number of experimentally identified exonic enhancers. Unfortunately, such a dataset is still lacking, because enhancers characterized by high throughput methods such as self-transcribing active regulatory regions sequencing (STARR-seq) [40] have up to 56% [41] and 87% [42] FDRs, respectively. Thus, a set of accurately predicted CRMs and constituent TFBSs might be an alternative solution to address these interesting and important questions about exonic CRMs.

We recently predicted a set of 1.41M CRM candidate (CRMCs) and a set of 1.96M non-CRMCs in the human genome by integrating more than 6,000 TF ChIP-seq datasets from various cell/tissue types [35]. These CRMCs harbor a total of Validation on experimentally determined CRM function-related sequence elements such as VISTA enhancers [43] and non-coding ClinVar variants [44] indicates that our predicted CRMCs are highly accurate, with an estimated FDR of 0.05% and false negative rate (FNR) of 1.27% [35]. The FDR and FNR of predicted CRMs can decrease further when a more stringent p-value cutoff is used [35]. As expected, most (∼94%) of the nucleotide positions of the predicted CRMs and constituent TFBSs are located in non-exonic sequences (NESs), while the remaining ∼6% overlap exons [35]. These predicted exonic CRMs and TFBSs promoted us to re-examine the duon hypothesis and address related fundamental questions for the dual-use of exons including CDSs and UTRs that have not been examined before.

## Results

### Exonic TFBSs are prevalent but unevenly distributed along CDSs and UTRs

To analyze potential exonic *cis*-regulatory sequences in the human genome and address the related issues, we used the 428,628 CRMs containing 38,507,543 putative TFBSs predicted at a stringent p-value cutoff of 5x10^-6^ to minimize FDR [35]. These CRMs and TFBSs have a total length of 982,470,181bp and 397,703,041bp, covering 31.81% and 12.88% of the mappable genome (3,088,269,832bp), respectively [35]. As expected, the vast majority (94.42%) of the CRM positions are located in NESs, while the remaining 5.58% fall in exons. There are a total of 180,223 (34.08%) CRMs that at least partially overlap exons, and we refer them as exonic CRMs. Only small number (0.16%) of exonic CRMs entirely fall in exons, most of them have less than 10% of their lengths overlapping exons (Figure 1a). Figures S1a∼S1k (Additional File 1) shows 11 examples of the exonic CRMs that overlap experimentally verified exonic enhancers. In these examples, some exonic CRMs overlap one or few exons and spans adjacent introns and/or intergenic regions ((Additional File 1, Figures S1a∼S1h), some others span the entire locus of a gene (Additional File 1, Figures S1i∼S1k).

**Figure 1.**
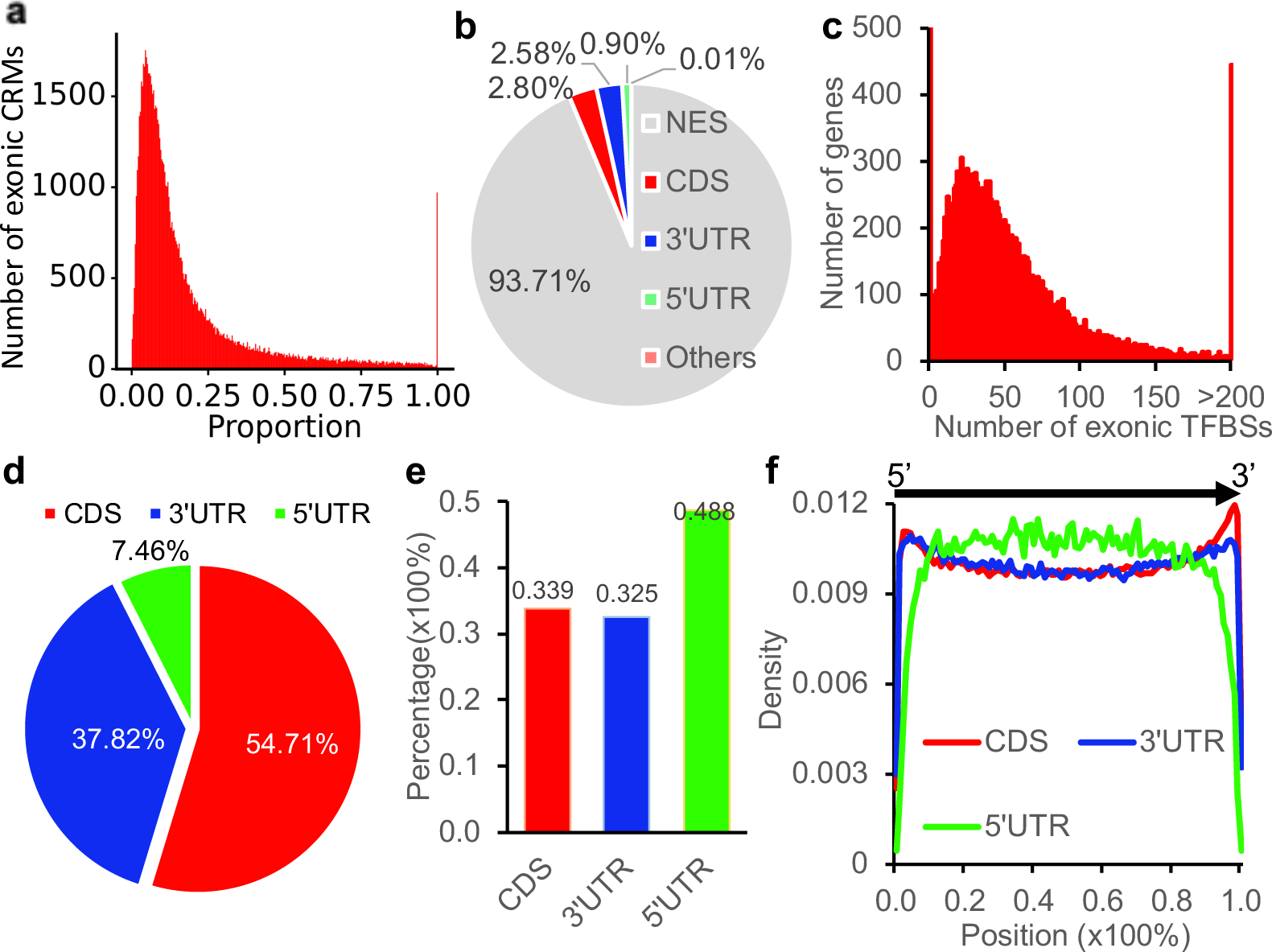
Properties of the predicted exonic CRMs and exonic TFBSs in the human genome. **a.** Number of exonic CRMs with different proportions of exonic sequences. **b.** Proportions of predicted TFBS positions falling in NESs, CDSs, 5’-UTRs, 3’-UTRs or other sequences (with both 5’-UTR and 3’-UTR annotations). **c.** Number of genes containing different numbers of predicted exonic TFBSs. **d**. Percentages of exonic TFBS positions falling in CDSs, 5’-UTRs or 3’-UTRs. **e**. Percentage of exonic TFBS positions in the total length of CDSs, 5’-UTRs or 3’-UTRs. **f**. Distribution of exonic TFBSs along CDSs, 5’-UTRs and 3’-UTRs of genes from the 5’-end to the 3’-end as indicated by the horizontal arrow. The CDS of a gene is made by concatenating all coding exons of along the gene.

The vast majority (93.71%) of TFBS positions are located in NESs, while the remaining 6.29% reside in exons (Figure 1b). We consider a TFBS as a non-exonic TFBS if it has at least one position located in a NES. Of the 38,507,543 putative TFBSs in the CRMs, 36,597,744 (95.04%) are non-exonic TFBSs, while the remaining 1,909,799 (4.96%) TFBSs are entirely located within an exon; we consider them as exonic TFBSs. We focus our subsequent analyses mainly on these exonic TFBSs instead of the exonic CRMs, since it is the constituent TFBSs in a CRM that mainly determine the function of the CRM [45, 46]. If an exonic TFBS position is annotated to be in both a CDS and a UTR, we consider it to be in the CDS. If an exonic TFBS position is annotated to be in both a 5’-UTR and a 3’-UTR, we consider it as “others” and exclude it from analyses. Of all the TFBS positions (397,703,041bp), 2.80%, 0.90%, 2.58% and 0.01% are located in CDSs, 5’-UTRs, 3’-UTRs and “others” respectively (Figure 1b). If an exonic TFBS is entirely located in a CDS, a 5’-UTR or a 3’-UTR, we refer it as a coding TFBS, a 5’-UTR TFBS or a 3’-UTR TFBS, respectively. Of the 38,507,543 putative TFBSs, 1,047,183 (2.72%) 129,518 (0.34%) and 733,089 (1.90%) are coding TFBSs, 5’-UTR TFBSs and 3’-UTR TFBSs, respectively. The exonic TFBSs are located in exons of 19,060 (96.78%) of the 19,694 transcriptionally verified genes, and each gene harbored an average of 56 potentially overlapping exonic TFBSs (Figure 1c). More specifically, of the exonic TFBS positions, 54.71%, 7.46% and 37.82% are located in CDSs, 5’-UTRs and 3’-UTRs of 18,971 (96.32%), 12,903 (65.52$) and 13,237 (67.21%) genes, respectively (Figure 1d). Thus, the vast majority of genes have at least a coding exon, 5’-UTR or 3’-UTR at least partially overlapping an exonic TFBS. The coding TFBSs, 5’-UTR TFBSs and 3’-UTR TFBSs comprise 33.9%, 48.8% and 32.5% of the total length of the annotated CDSs (32,797,669bp), 5’-UTRs (3,104,596bp) and 3’-UTRs (23,630,714bp), respectively (Figure 1e). Therefore, a third of CDS and 3’-UTR lengths are exonic TFBS positions, while almost a half of 5’-UTR lengths are exonic TFBS positions, indicating that exonic TFBSs as well as exonic CRMs might be more prevalent than originally thought [30, 47]. Interestingly, coding TFBS and 3’-UTR TFBS positions tend to be clustered at the two ends of CDSs and 3’-UTRs, respectively, while 5’-UTR TFBS positions are evenly located along most part of 5’-UTRs but tend to avoid the two ends (Figure 1f).

### C/G contents elevate at the degenerate third codon positions in coding TFBSs as well as in 5’-UTR TFBSs and non-exonic TFBSs

To evaluate possible biased use of A/T and C/G in our predicted TFBSs and non-CRMCs, we first compared nucleotide frequencies at the degenerate third codon positions of the 21 synonymous codon sets in coding TFBSs and non-CRMC CDSs (for the definition of the degenerate third codon positions, see Materials and Methods). As shown in Figure 2a, although the human genome is A/T-rich (59% A/T vs 41% C/G) [48], in coding TFBSs, there are more C/G than A/T at the degenerate third codon positions of 17 of the 21 synonymous codon sets, except for Asn (AA[CT]), Ile (AT[ACT]), Ser (TC*) and Thr (AC*). In contrast, in non-CRMC CDSs, there are more A/T than C/G at the degenerate third codon positions of 17 of the 21 synonymous codon sets, except for Gln (CA[AG]), Leu (CT*), Leu (TT[AG]) and Val (GT*) (Figure 2a). We next compared nucleotide frequencies at the degenerate third codon positions of the 21 synonymous codon sets in coding TFBSs relative to those in non-CRMC CDSs. As shown in Figure 2b, the log odd ratios of the frequencies indicate that C/G are more preferred at the degenerate third codon positions in coding TFBSs than at those in non-CRMC CDSs (p< 2.2x10^-16^, χ^2^ test) for all the 21 synonymous codon sets. This result is in agreement with the earlier finding that C/G was more preferred at the degenerate third codon positions in the 21 synonymous codon sets in duons than those in non-duon CDSs [33].

**Figure 2.**
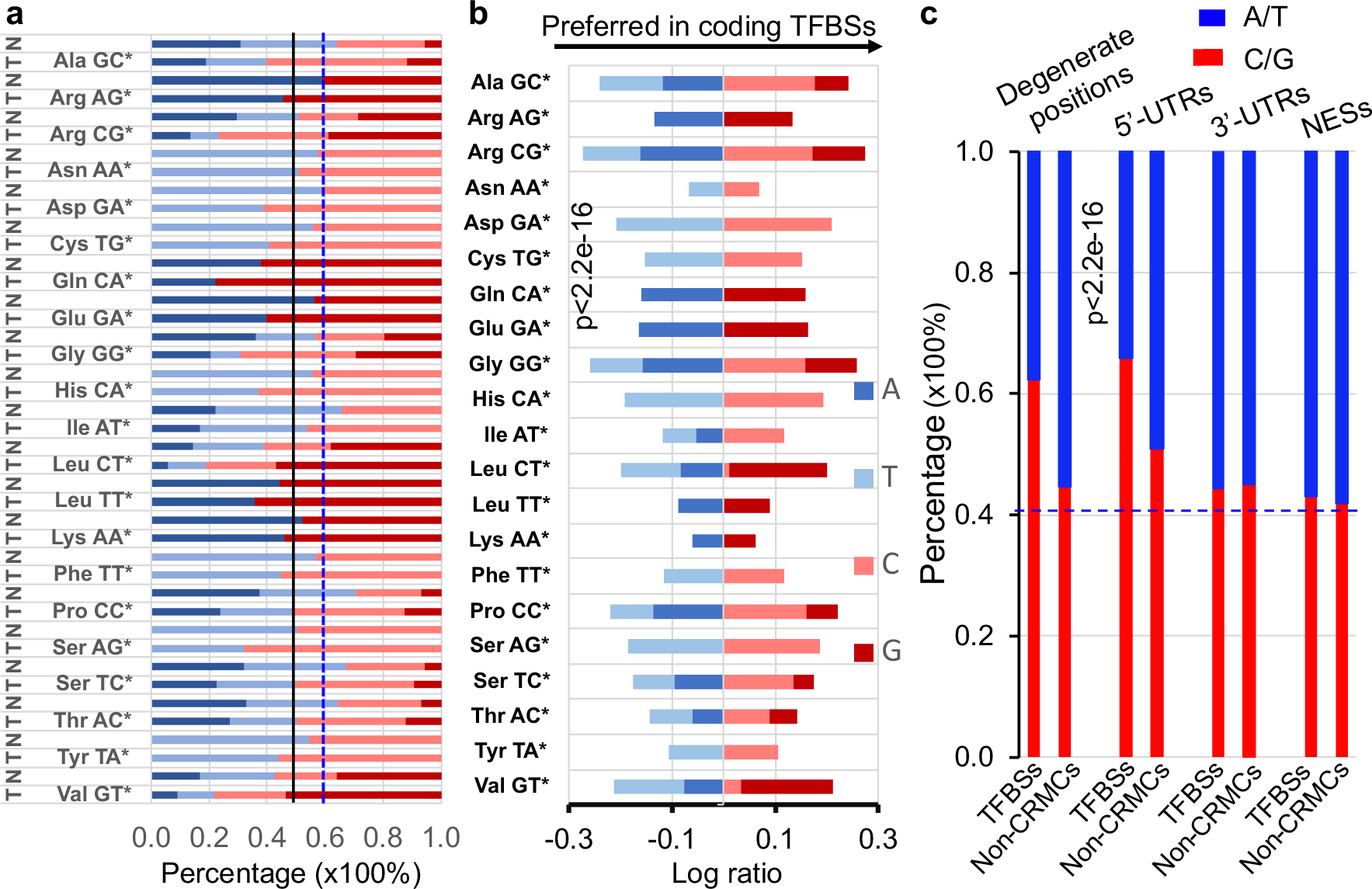
Biased distribution of A/T and C/G in predicted TFBSs and non-CRMCs. **a.** Nucleotide frequencies at the degenerate third positions of the 21 synonymous codon sets in coding TFBSs (T) and in non-CRMC CDSs (N). The dotted vertical line indicates the neutral expectation (59%) of A/T contents in the genome. **b.** Log ratio of A/T and C/G contents at the degenerate third codon positions of the 21 synonymous codon sets in coding TFBSs over the those in non-CRMC CDSs. P<2.2x10^-16^ for each codon set was computed using the χ^2^ test. **c.** A/T and C/G contents in the degenerate third codon positions, 5’-UTRs, 3’-UTRs and NESs in TFBSs in comparison with those in their counterparts in non-CRMCs. The dotted horizontal line indicates the neutral expectation (41%) of C/G contents in the genome. P<2.2x10^-16^ for each comparison was computed using the χ^2^ test.

We also compared A/T and C/G contents at the degenerate third codon positions of in coding TFBSs, at 5’-UTR TFBSs, at 3’-UTR TFBSs and at non-exonic TFBSs with those in their non-CRMC counterparts. As expected, C/G contents (62.29%) at the degenerate third codon positions in coding TFBSs are much higher than those (44.62%) in non-CRMC CDSs (Figure 2c, p< 2.2x10^-16^, χ^2^ test). Interestingly, 5’-UTR TFBSs (65.77%) and non-exonic TFBSs (43.08%) also have significantly higher C/G contents than do non-CRMC 5’-UTRs (50.85%) (Figure 2c, p< 2.2x10^-16^, χ^2^ test) and non-CRMC NESs (41.90%) (Figure 2c, p< 2.2x10^-16^, χ^2^ test), respectively. However, 3’-UTR TFBSs (44.23%) have slightly but significantly lower C/G contents than non-CRMC 3’-UTRs (44.96%) (Figure 2c, p< 2.2x10^-16^, χ^2^ test). Moreover, C/G contents in the degenerate third codon positions in non-CRMC CDSs (44.62%), non-CRMC 5’-UTRs (50.85%) and 3’-UTRs (44.96%) are also higher (Figure 2c, p< 2.2x10^-16^, χ^2^ test) than the neutral expectation (41%) in the genome (Figure 2c). It has been shown that C/G contents at four-fold degenerate positions are consistently elevated above the neutral expectation in all genomes examined [49]. Thus, with C/G contents of 62.29% (Figure 2c), this is particularly true for degenerate third codon positions in coding TFBSs. Interestingly, while non-CRMC NESs have the similar C/G contents (41.90%) to the neutral expectation (41%), non-exonic TFBSs (43.08%) have higher C/G contents than the neutral expectation (41%) (Figure 2c). It is believed that the elevation of C/G contents in genome regions is caused by G/C-biased gene conversion during miotic recombination and DNA repairing [50]. As nucleosome-free DNA is more susceptible to damage than nucleosome wrapped DNA [51], the elevation of C/G contents in coding TFBSs (62.29%) and 5’-UTR TFBSs (65.77%) might be due to their prolonged nucleosome-free states during transcription and TF binding compared to their non-CRMC counterparts; while the elevation of C/G contents (44.62%) in non-exonic TFBS positions might be due to their prolonged nucleosome-free states during TF binding compared to non-CRMC NESs (41.90%). The much higher C/G contents (62.29%) at the degenerate third codon positions in coding TFBSs and 5’-UTR TFBSs (65.77%) than in non-exonic TFBS positions (43.08%) might be due to the possible longer time that coding TFBSs and 5’-UTR TFBSs are in the nucleosome-free state than are non-exonic TFBSs. The slightly higher C/G contents (44.96%) in non-CRMC 3’-UTRs than in 3’-UTR TFBSs (44.23%) (Figure 2c) suggest that some 3’-UTR related functions carried out by the middle part of 3’-UTRs where 3’-UTR TFBSs are depleted (Figure 1f) might spend more time in the nucleosome-free state than do 3’-UTR TFBSs.

### Non-degenerate first and second codon positions as well as degenerate third codon positions in coding TFBSs are more likely under evolutionary constraints than are those in non-CRMC CDSs

Since the two earlier studies [30, 33] drew contradictory results about the evolution of duons predicted based on DHSs it is interesting to see how our predicted coding TFBSs evolve when compared with non-CRMC CDSs. The highly biased C/G contents in TFBSs versus in non-CRMCs prompted us to consider A/T and C/G positions separately to avoid pitfalls due to different evolutionary behaviors of A/T and C/G in the genome [33]. To this end, we first compared the evolutionary profiles of A/T and C/G at the non-degenerate first and second codon positions as well as the degenerate third codon positions (for the definitions of non-degenerate and degenerate codon positions, see Materials and Methods) in coding TFBSs with those in non-CRMC CDSs. We used phyloP scores [32] of nucleotide positions in the human genome for this purpose, since phyloP treats negative and positive selections in a unified manner and detects departures from the neutral rate of substitution in either direction, while allowing for clade-specific selection. Also, phyloP scores are available for the vast majority of positions in the human genome (https://genome.ucsc.edu/), making our analysis more robust. The larger a positive phyloP score of a nucleotide position, the more likely it is under negative selection; the smaller a negative phyloP score of a position, the more likely it is under positive selection; and a phyloP score around zero means that the position is selectively neutral or nearly so [32]. For the convenience of discussion, we consider a nucleotide position with a phyloP score within an interval around 0 [−δ, δ] (δ > 0) to be selectively neutral or nearly so, a position with a score greater than δ to be under negative selection, and a position with a score less than – δ to be under positive selection. We define the size of the area under the density curve of the distribution of the scores of a set of genomic positions within the intervals [−δ, δ], (δ, *maximum*) and (*minimum* −δ), to be the proportion of the positions that are selectively neutral or nearly so, under negative selection and under positive selection, respectively. In this study, we chose δ =1.3, corresponding to a p-value of 0.05 to claim a position to be under negative or positive selection [32]. It is reasonable to assume that evolutionary constraints on the non-degenerate first and second codon positions reflect selection pressures both for encoding amino acids and other functions if any, such as TF binding; while evolutionary constraints on the degenerate third codon positions indicate selection pressures for other functions such as TF binding, other than coding specific amino acids

Both A/T and C/G at both the non-degenerate first (Figure 3a) and second (Figure 3b) codon positions in coding TFBSs have significantly different phyloP score distributions than those in non-CRMC CDSs (p<2.23x10^-302^, K-S test), suggesting that the non-degenerate positions in coding TFBSs evolve differently from those in non-CRMC CDSs. Specifically, as expected, most (>60%) of the non-degenerate first (Figure 3c) and second (Figure 3d) codon positions are under negative selection, regardless of where they are located, i.e., in coding TFBSs or in non-CRMC CDSs. However, there are higher proportions of A/T (70.51% vs 60.91%, difference: 9.6%, p < 2.2x10^-16^, χ^2^ test) and C/G (68.92% vs 62.67%, difference: 6.25%, p < 2.2x10^-16^, χ^2^ test) at the non-degenerate first codon positions in coding TFBSs to be under negative selection than those in non-CRMC CDSs (Figure 3c). There are also higher proportions of A/T (78.52% vs 70.92%, difference: 7.6%, p < 2.2x10^-16^, χ^2^ test) and C/G (69.70% vs 63.05%, difference: 6.65%, p < 2.2x10^-16^, χ^2^ test) at the non-degenerate second codon positions in coding TFBSs to be under negative selection than those in non-CRMC CDSs (Figure 3d). Only a small proportion (∼3.5%) of non-degenerate first (Figure 3c) and second (Figure 3d) positions are under positive selection, regardless of where they are located, i.e., in coding TFBSs or in non-CRMC CDSs. However, there are slightly lower proportions of A/T (2.76% vs 3.46%, difference: -0.7%, p < 2.2x10^-16^, χ^2^ test) and C/G (2.28% vs 3.33%, difference: -1.05%, p < 2.2x10^-16^, χ^2^ test) at the non-degenerate first codon positions in coding TFBSs to be under positive selection than those in non-CRMC CDSs (Figures 3c). There are also slightly lower proportions A/T (1.82% vs 2.20%, difference: -0.38%, p < 2.2x10^-16^, χ^2^ test) and C/G (2.28% vs 3.71%, difference: -1.43%, p < 2.2x10^-16^, χ^2^ test) at the non-degenerate second codon positions in coding TFBSs to be under positive selection than those in non-CRMC CDSs (Figure 3d). Consequently, up to slightly more than third of the non-degenerate first (Figure 3c) and second (Figure 3d) codon positions are selectively neutral or nearly so, regardless of where they are located, i.e., in coding TFBSs or in non-CRMC CDSs. However, there are lower proportions of A/T (26.74% vs 35.63%, difference: -8.89%, p < 2.2x10^-16^, χ^2^ test) and C/G (28.80% vs 34.00%, difference: -5.20%, p < 2.2x10^-16^, χ^2^ test) at the non-degenerate first codon positions in coding TFBSs to be selectively neutral or nearly so than those in non-CRMC CDSs (Figure 3c). There are also lower proportions of A/T (19.65% vs 26.88%, difference: -7.23%, p< 2.2x10^-16^, χ^2^ test) and C/G (28.02% vs 33.24%, difference: -5.22%, p < 2.2x10^-16^, χ^2^ test) at the non-degenerate second codon positions in coding TFBSs to be selectively neutral or nearly so than those in non-CRMC CDSs (Figure 3d). The differences between the proportions of positions that are selectively neutral in coding TFBSs and non-CRMC CDSs suggests that 8.89% and 7.23% of A/T and 5.20% and 5.22% of C/G at the non-degenerate first and second codon positions in the coding TFBSs, respectively, which are mostly under negative selections, might be involved in TF-DNA interactions. In other words, these positions might be in dual-use, i.e., for encoding amino acids in proteins and TF binding interfaces in DNA.

**Figure 3.**
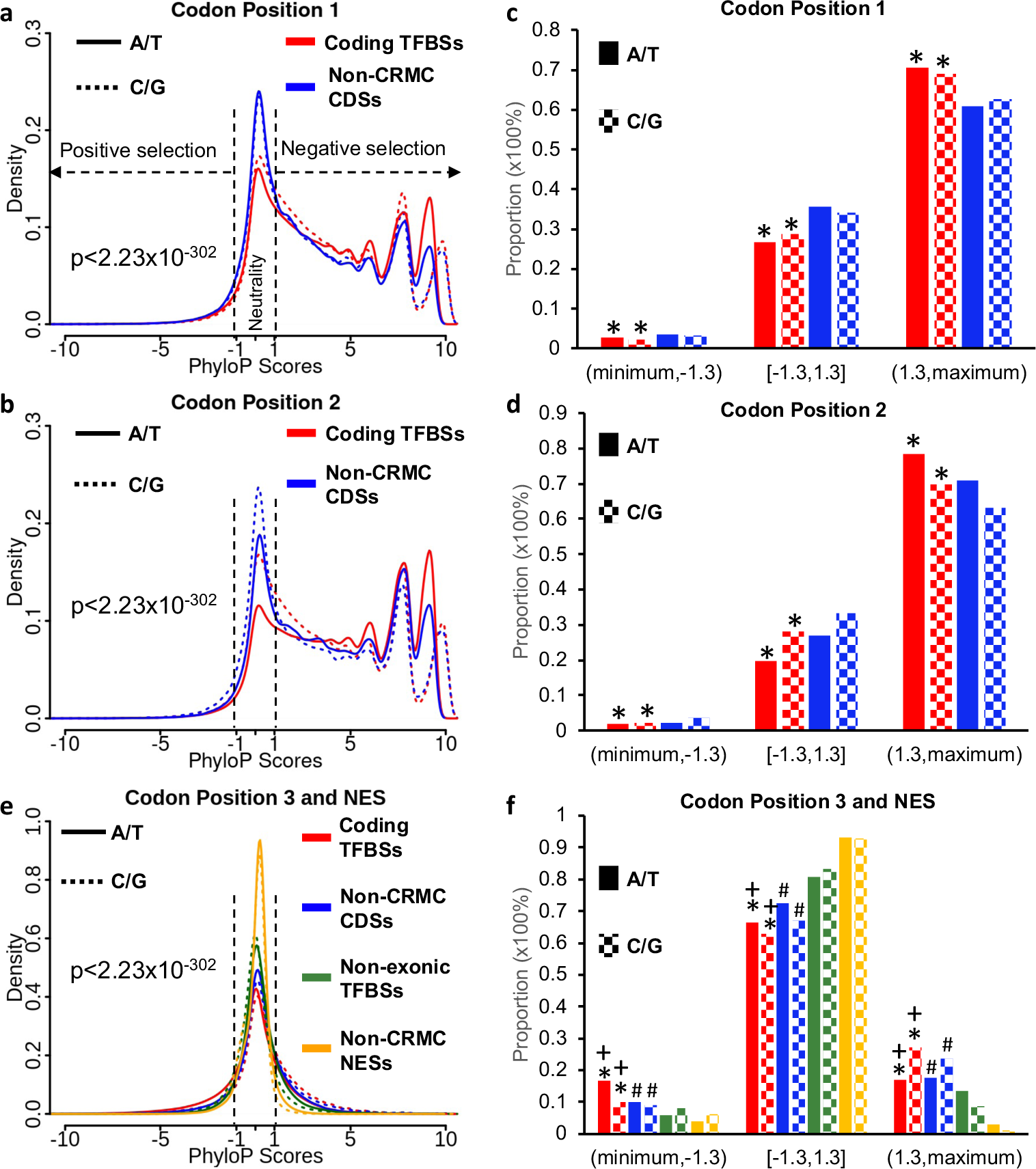
Comparison of phyloP scores of the TFBS positions in different types of sequences. **a, b, e**. Distributions of phyloP scores of A/T and C/G at the nondegenerate first codon positions in coding TFBSs and non-CRMC CDSs (**a**), at the nondegenerate second codon positions in coding TFBSs and non-CRMC CDSs (**b**), and at the degenerate third codon positions in coding TFBSs and non-CRMC CDSs, in comparison with those at non-exonic TFBS and non-CRMC NES positions (**e**). Comparisons of density curves were done using the K-S test. **c, d, f**. Proportions of A/T and C/G under positive selection [(minimum, -1.3)], negative selection [(1.3, maximum)], or of selective neutrality [[-1.3, 1.3]] at the nondegenerate first codon positions in coding TFBSs and non-CRMC CDSs (**c**), at the nondegenerate second codon positions in coding TFBSs and non-CRMC CDSs (**d**), and at the degenerate third codon positions in coding TFBSs and non-CRMC CDSs, in comparison with those at non-exonic TFBS and non-CRMC NES positions (**f**). *P < 2.2x10^-16^, comparisons between codon positions in coding TFBSs (red) and in non-CRMC CDSs (blue) using the χ^2^ test. +P < 2.2x10^-16^, comparisons between degenerate third codon positions in coding TFBSs (red) and non-exonic TFBS positions (green) using the χ^2^ test. #P < 2.2x10^-16^, comparisons between degenerate third codon positions in non-CRMC CDSs (blue) and NES positions (yellow) using the χ^2^ test.

Both A/T and C/G at the degenerate third codon positions (Figure 3e) in coding TFBSs also have significantly different phyloP score distributions than those in non-CRMC CDSs (p<2.23x10^-302^, K-S test), suggesting that these degenerate positions in coding TFBSs evolve differently from those in non-CRMC CDSs. As expected, in stark contrast to the non-degenerate first and second codon positions with >60% under negative selection (Figures 3c and 3d), less than 30% of degenerate third codon positions are under negative selection, regardless of where they are located, i.e., in coding TFBSs or in non-CRMC CDSs (Figure 3f). However, C/G (27.08%) at the degenerate third codon positions in coding TFBSs have a higher proportion to be under negative selection than those (23.72%) in non-CRMC CDSs (difference: 3.36%, p < 2.2x10^-16^, χ^2^ test) (Figure 3f), while the opposite (difference: -0.74%, p < 2.2x10^-16^, χ^2^ test) is true for A/T at the degenerate third codon positions in coding TFBSs (16.88%) and non-CRMC CDSs (17.62%) (Figures 3f). Interestingly, A/T at the degenerate third codon positions in coding TFBSs have a higher proportion (16.59%) to be under positive selection than those (9.94%) in non-CRMC CDSs (difference: 6.65%, p < 2.2x10^-16^, χ^2^ test) (Figure 3f). C/G at the degenerate third codon positions in coding TFBSs also have a slightly higher proportion (10.07%) to be under positive selection than those (9.12%) in non-CRMC CDSs (difference: 0.95%, p < 2.2x10^-16^, χ^2^ test) (Figure 3f). Consequently, both A/T (65.53%) and C/G (62.85%) at the degenerate third codon positions in coding TFBSs have a lower proportion of selective neutrality than those (72.44% and 67.16%, respectively) in non-CRMC CDSs (differences: -6.91% and -4.31%, respectively, p<2.2x10^-16^, χ^2^ test). The difference in the proportions of positions that are selectively neutral between coding TFBSs and non-CRMC CDSs suggests that about 6.91% of A/T and 4.31% of C/G at the degenerate third codon positions in coding TFBSs, which are either under positive (particularly, for A/T) or negative (particularly, for C/G) selections, might be involved in TF-DNA interactions. This conclusion therefore is in sharp contrast to either of the earlier two that the degenerate third codon positions in duons are more conserved than those in non-duons [30], or that they are similarly or only slightly more conserved than those in non-duon CDSs [33]. These results indicate that our predicted coding TFBSs are distinct from the duons used in the earlier two studies, in which the duons were predicted based on DHSs and thus might not necessarily be TFBSs or parts of CRMs. Taken together, these results suggest that a considerable proportion of the non-degenerate first (8.89% of A/T and 5.20% of C/G) and second (7.23% of A/T and 5.22% of C/G) codon positions as well as the degenerate third (6.91% of A/T and 4.31% of C/G) codon positions in coding TFBSs might be involved in TF-DNA interactions for transcriptional regulation.

### Degenerate third codon positions in coding TFBSs are more likely under either positive or negative selections than are non-exonic TFBSs

We reason that if the degenerate third codon positions in coding TFBSs are only for TF binding other than being a wobble base in the codon, then they should evolve similarly to those in non-exonic TFBSs that are entirely located in NESs, and thus for at least most of them, the main functions might be for TF binding. However, as shown in Figure 3f, both A/T (16.88%) and C/G (27.08%) at the degenerate third codon positions in coding TFBSs have a higher proportion to be under negative selection than those (13.34% and 8.68%, respectively) in non-exonic TFBS positions (difference: 3.54% and 18.40%, respectively, p < 2.2x10^-16^, χ^2^ test). Moreover, both A/T (16.59%) and C/G (10.07%) at the degenerate third codon positions in coding TFBSs also have a higher proportion to be under positive selection than those (5.89% and 8.22%, respectively) at non-exonic TFBS positions (difference: 10.70% and 1.85%, respectively, p < 2.2x10^-16^, χ^2^ test) (Figure 3f). Consequently, both A/T (66.53%) and C/G (62.85%) at the degenerate third codon positions in coding TFBSs have a lower proportion of selective neutrality than those (80.77% and 83.10%, respectively) at non-exonic TFBS positions (difference: -14.24% and -20.25%, respectively, p < 2.2x10^-16^, χ^2^ test) (Figure 3f). These results suggest that in addition to TF binding, 14.24% of A/T and 20.25% of C/G at the degenerate third codon positions in coding TFBSs might have other functions such as splicing regulation [52–54], and thus, might be in dual-use.

### Degenerate third codon positions in non-CRMC CDSs are more likely under either positive or negative selections than are positions in non-CRMC NESs

Moreover, we reason that if the degenerate third codon positions in non-CRMC CDSs do not have any function except for being the wobble bases in codons, then they should evolve similarly to positions of non-CRMC NESs that are largely selective neutral [35]. However, as shown in Figure 3f, both A/T (17.62%) and C/G (23.72%) at the degenerate third codon positions in non-CRMC CDSs have a higher proportion to be under negative selection than those (3.09% and 1.23%, respectively) at non-CRMC NES positions (difference: 14.53% and 22.49%, respectively, p < 2.2x10^-16^, χ^2^ test). Moreover, both A/T (9.94%) and C/G (9.12%) at the degenerate third codon positions in non-CRMC CDSs also have a higher proportion to be under positive selection than those (3.80% and 6.03%, respectively) at non-CRMC NES positions (difference: 6.14% and 3.09%, respectively, p < 2.2x10^-16^, χ^2^ test) (Figure 3f). Consequently, both A/T (72.44%) and C/G (67.16%) at the degenerate third codon positions in non-CRMC CDSs have a lower proportion of selective neutrality than those (93.11% and 92.73%, respectively) at non-CRMC NES positions (difference: -20.67% and -25.57%, respectively, p < 2.2x10^-16^, χ^2^ test) (Figure 3f). These results suggest that 20.67% of A/T and 25.57% of C/G at the degenerate third codon positions in non-CRMC CDSs might have other biological functions such as splicing regulation [52–54], in addition to being wobble bases.

### 5’-UTR TFBSs and 3’-UTR TFBSs are more likely under evolutionary constraints than are corresponding non-CRMC UTRs

To assess how 5’-UTR TFBSs and 3’-UTR TFBSs evolve, we compared the phyloP scores of their A/T and G/C positions with those in their non-CRMC counterparts. As shown in Figure 4a, both A/T and C/G positions at 5’-UTR TFBSs have significantly different (p<2.23x10^-302^, K-S test) phyloP score distributions than those in non-CRMC 5’-UTRs, suggesting that 5’-UTR TFBS positions evolve differently from non-CRMC 5’-UTR positions. Specifically, both A/T (17.81%) and C/G (20.88%) positions in 5’-UTR TFBSs have a higher proportion to be under negative selection than those (5.90% and 4.54%, respectively) in non-CRMC 5’-UTRs (differences: 11.91% and 16.34%, respectively, p < 2.2x10^-16^, χ^2^ test) (Figure 4b). A/T positions (9.73%) in 5’-UTR TFBSs have a higher proportion under positive selection than those (5.89%) in non-CRMC 5’-UTRs (differences: 3.84%, p < 2.2x10^-16^, χ^2^ test), but the opposite is true for C/G positions (difference: -1.72%, Figure 4b). Consequently, both A/T (72.45%) and C/G (72.62%) positions in 5’-UTR TFBSs have a lower proportion of selective neutrality than those (88.31% and 88.78%, respectively) in non-CRMC 5’-UTRs (differences: -15.68% and -16.16%, respectively, p < 2.2x10^-16^, χ^2^ test) (Figure 4b). These results suggest that about 15.68% of A/T and 16.16% C/G positions in 5’-UTR TFBSs that are under either positive or negative selections might be involved in TF-DNA interactions.

**Figure 4.**
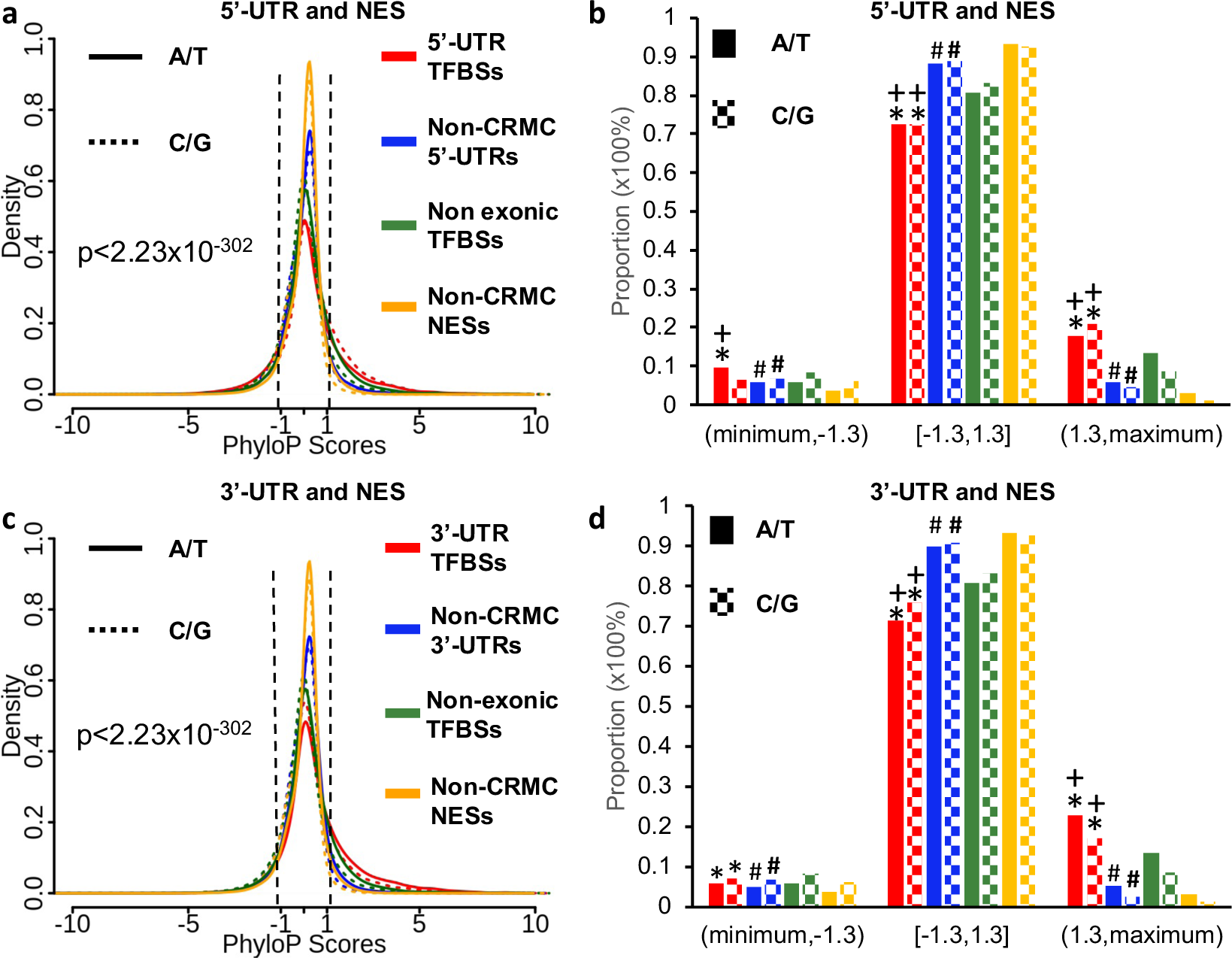
Comparison of phyloP scores of the UTR TFBS, non-CRMC UTR, non-exonic TFBS and non-CRMC NES positions. **a, c.** Distributions of phyloP scores of A/T and C/G positions in 5’-UTR TFBSs and non-CRMC 5’-UTRs (**a**), and in 3’-UTR TFBSs and non-CRMC 3’-UTRs (**c**), in comparison with those in non-exonic TFBSs and non-CRMC NESs. Comparisons of density curves were done using the K-S test. **b, d**: Proportions of the positions that are under positive selection [(minimum, -1.3)], negative selection [(1.3, maximum)], or of selective neutrality [[-1.3, 1.3])], in 5’-UTR TFBSs and non-CRMC 5’-UTRs (**c**), and in 3’-UTR TFBSs and non-CRMC 3’-UTRs (**d**), in comparison with those in non-exonic TFBSs and non-CRMC NESs. *P < 2.2x10^-16^, comparisons between UTR TFBS positions (red) and non-CRMC UTR positions (blue) using the χ^2^ test. +P < 2.2x10^-16^, comparisons between UTR TFBS positions (red) and non-exonic TFBS positions (green) using the χ^2^ test. #P < 2.2x10^-16^, comparisons between non-CRMC UTR positions (blue) and NES positions (yellow) using the χ^2^ test.

Moreover, as shown in Figure 4c, both A/T and C/G positions in 3’-UTR TFBSs also have significantly different (p<2.23x10^-302^, K-S test) phyloP score distributions than those in non-CRMC 3’-UTRs, suggesting that 3’-UTR TFBS positions evolve differently from non-CRMC 3’-UTR positions. Specifically, both A/T (22.93%) and C/G (17.15%) positions in 3’-UTR TFBSs have a higher proportion to be under negative selection than those (5.16% and 2.40%, respectively) in non-CRMC 3’-UTRs (differences: 17.77% and 14.75%, respectively, p < 2.2x10^-16^, χ^2^ test) (Figure 4d). Both A/T (5.74%) and C/G (6.97%) positions in 3’-UTR TFBSs have a slightly higher proportion to be under positive selection than those (5.01% and 6.75, respectively) in non-CRMC 3’-UTRs (differences: 0.73% and 0.22% p < 2.2x10^-16^, χ^2^ test) (Figure 4d). Consequently, both A/T (71.34%) and C/G (75.88%) positions in 3’-UTR TFBSs have a lower proportion of selective neutrality than those (89.83% and 90.85%, respectively) in non-CRMC 3’-UTRs (differences: -18.49% and -14.97%, respectively, p < 2.2x10^-16^, χ^2^ test) (Figure 4d). These results suggest that about 18.49% of A/T and 14.97% C/G positions in 3’-UTR TFBSs that are under either positive or negative selection might be involved in TF-DNA interactions.

### 5’-UTR TFBSs and 3’-UTR TFBSs are more likely under evolutionary constraints than are non-exonic TFBSs

We reason that if 5’-UTR TFBS and 3’-UTR TFBS positions are only for TF binding, then they should evolve similarly to those in non-exonic TFBSs that are entirely located in NESs.

However, as shown in Figure 4d, both A/T (17.81%) and C/G (20.88%) in 5’-UTR TFBSs have higher proportions of positions under negative selection than those (13.34% and 8.68%, respectively) in non-exonic TFBSs (differences: 4.47% and 12.20%, respectively). A/T in 5’-UTR TFBSs have a higher proportion (9.73%) of positions under positive selection than those (5.89%) in non-exonic TFBSs (differences: 3.84%, p < 2.2x10^-16^, χ^2^ test), but the opposite is true for C/G (difference: -1.72%, Figure 4d). Consequently, both A/T (72.45%) and C/G (72.62%) in 5’-UTR TFBSs have a lower proportion of positions that are selectively neutral than those (80.77% and 83.10%, respectively) in non-exonic TFBSs (differences: -8.32% and -10.48%, respectively, p < 2.2x10^-16^, χ^2^ test) (Figure 4b). These results suggest that about 8.32% of A/T and 10.48% C/G in 5’-UTR TFBSs that are under either positive or negative selections might have other 5’-UTR-related functions in addition to TF binding, and thus, might be in dual-use.

Moreover, as shown in Figure 4d, both A/T (22.93%) and C/G (17.15%) positions in 3’-UTR TFBSs have higher proportions to be under negative selection than those (13.34% and 8.68%, respectively) in non-exonic TFBSs that are entirely in NESs (differences: 9.59% and 8.47%, respectively, p < 2.2x10^-16^, χ^2^ test). Moreover, both A/T (5.74%) and C/G (6.97%) positions in 3’-UTR TFBSs have lower proportions to be under positive selection than those (5.89% and 8.22%, respectively) in non-exonic TFBSs that are entirely in NESs (differences: - 0.15% and -1.25%, respectively) (Figure 4d). Nonetheless, both A/T (71.34%) and C/G (75.88%) positions in 3’-UTR TFBSs have a lower proportion of selective neutrality than those (80.77% and 83.10%, respectively) in non-exonic TFBSs (differences: -9.43% and -7.22%, respectively, p < 2.2x10^-16^, χ^2^ test) (Figure 4d). These results suggest that about 9.43% of A/T and 7.22% of C/G positions in 3’-UTR TFBSs that are under either positive or negative selection might have other 3’-UTR-related functions in addition to TF binding, and thus, might be in dual-use.

### Non-CRMC 5’-UTRs and non-CRMC 3’-UTRs have higher proportions of positions under either negative selection or positive selection than non-CRMC NESs

We also reason that if Non-CRMC 5’-UTR and non-CRMC 3’-UTR positions do not have any function, then they should evolve similarly to non-CRMC NES positions that are largely selective neutral [35]. However, as shown in Figure 4b, both A/T (5.90%) and C/G (4.54%) positions in non-CRMC 5’-UTRs have higher proportions to be under negative selection than those (3.09% and 1.23%, respectively) in non-CRMC NESs (differences: 2.81% and 3.31%, respectively, p < 2.2x10^-16^, χ^2^ test). Both A/T (5.80%) and C/G (6.68%) positions in non-CRMC 5’-UTRs have higher proportions of positions under positive selection than those (3.80%% and 6.03%, respectively) in non-CRMC NESs (differences: 2.00% and 0.65%, respectively, p < 2.2x10^-16^, χ^2^ test) (Figure 4c). Consequently, both A/T (88.31%) and C/G (88.78%) positions in non-CRMC 5’-UTRs have lower proportions of selective neutrality than those (93.11%% and 92.73%, respectively) in non-CRMC NESs (differences: -4.8% and -3.95%, respectively, p < 2.2x10^-16^, χ^2^ test) (Figure 4c). These results suggest that about 4.8% of A/T and 3.95% of C/G positions in non-CRMC 5’-UTRs, might be involved in 5’-UTR-related functions, such as ribosome binding [55], which is consistent with the current understanding about the functions of 5’-UTRs [55].

Moreover, as shown in Figure 4d, both A/T (5.16%) and C/G (2.40%) positions in non-CRMC 3’-UTRs have higher proportions of positions to be under negative selection than those (3.09% and 1.23%, respectively) in non-CRMC NESs (differences: 2.07% and 1.17%, respectively, p < 2.2x10^-16^, χ^2^ test). Both A/T (5.01%) and C/G (6.75%) positions in non-CRMC 3’-UTRs have higher proportions to be under positive selection than those (3.80% and 6.03%, respectively) in non-CRMC NESs (differences: 1.20% and 0.72%, respectively, p < 2.2x10^-16^, χ^2^ test) (Figure 4d). Consequently, both A/T (89.83%) and C/G (90.85%) positions in non-CRMC 3’-UTRs have higher proportions of selective neutrality than those (93.11% and 92.73%, respectively) in non-CRMC NESs (differences: -3.18% and -1.88%, respectively, p < 2.2x10^-16^, χ^2^ test) (Figure 4d). These results suggest that about 3.18% of A/T and 1.88% of C/G positions in non-CRMC 3’-UTRs might be involved in 3’-UTR-related functions, such as RNA-binding protein (RBPs) binding [56], which is consistent with the current understanding about the functions of 3’-UTRs [56].

### Coding TFBSs preferentially code for loops rather than helices and strands

One of the puzzles for the dual-use of codons is how the two irrelevant functions of a DNA sequence can be possibly co-evolved. To address this, we mapped amino acids encoded by coding TFBSs to known 3D structures of proteins in protein data bank (PDB) (https://www.rcsb.org) (Materials and Methods). To reduce the biased distribution of structures to some protein families, we generated a non-redundant protein set with a 30% sequence identity cutoff (Materials and Methods), whose CDSs contained a total of 7,266,384bp coding TFBS positions. Amino acids encoded by 7.49% of the 7,266,384bp coding TFBS positions can be mapped to 1,761 known protein structures. These mapped amino acids are enriched in loops (48.73%, χ^2^ test) compared with the proportion of length of loops in host proteins (46.65%, p < 2.2x10^-16^) as well as in a non-redundant set of all known protein structures [57] (47.10%, p < 2.2x10^-16^, χ^2^ test) (Figure 5a). On the other hand, these mapped amino acids are under-represented in helices (33.53%) and strands (17.74%) compared with the proportions of helices and strands in the host proteins (34.31% and 19.04%, respectively, p < 2.2x10^-16^, χ^2^ test) as well as in all proteins with known structures [57] (34.21% and 18.69%, respectively, p < 2.2x10^-16^, χ^2^ test) (Figure 5a). For amino acids encoded by the remaining 92.51% of the coding TFBSs positions, which could not be mapped to any known protein structures, we predicted their secondary structure types using RaptorX [58]. A similar pattern is found showing that amino acids encoded by these coding TFBSs are enriched in the putative loops but depleted in the putative helices and strands (Figure 5b). Specifically, 60.79%, 30.01% and 9.20% of the peptides encoded by coding TFBSs are predicted to adopt loops, helices and strands, respectively, while these proportions are 57.30%, 31.92% and 10.78% (p < 2.2x10^-16^, χ^2^ test), respectively, for the host proteins; and 58.20%, 31.44% and 10.36% (p < 2.2x10^-16^, χ^2^ test), respectively, for all the proteins with predicted secondary structures (Figure 5b). It has been shown that loops are generally less conserved than helices and strands [57], and we see the same results for both known (Figure 5c) and predicted (Figure 5d) secondary structure types. Since the folding of a protein is mainly determined by its helix and strand structures, but not its loops, changes in amino acids in the loops are less likely to alter the overall folding and functions of the protein. In this regard, it is not surprising that coding TFBSs tend to encode amino acids in loops where negative selection are weaker (Figures 5c, 5d), and therefore they could adopt for specific TF binding without compromising the overall structures of proteins.

**Figure 5.**
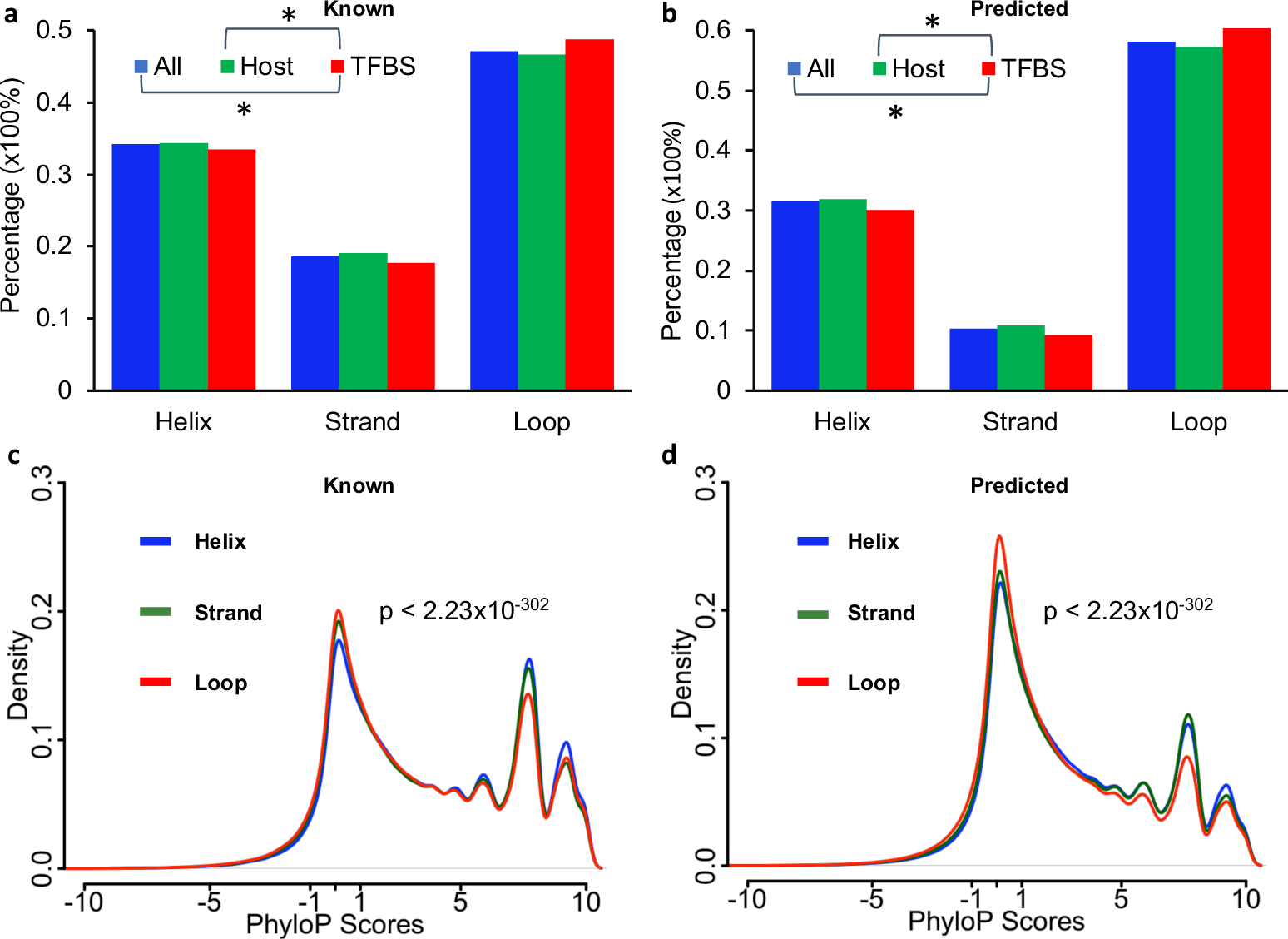
Preference of coding TFBSs for secondary structural types in proteins. **a.** Proportions of coding TFBS-encoded amino acids mapped to known loops, helices, and strands in comparison with those in the host proteins and in all known non-redundant protein structures. *P<2.2x10^-16^ was computed using the χ^2^ test for all the comparisons. **b.** Proportions of coding TFBS-encoded amino acids mapped to the predicted loops, helices, and strands in comparison with those in the host proteins and in all predicted secondary structures. *P<2.2x10^-16^ were computed using the χ^2^ test for all comparisons**. c.** Distributions of phyloP scores of coding TFBSs that encode amino acids mapped to known loops, helices, and strands. **d.** Distributions of phyloP scores of coding TFBSs that encode amino acids mapped to predicted loops, helices, and strands. P values (**c**) and (**d**) were computed using the K-S test.

### Active exonic TFBSs tend to be in close physical proximity to distal promotors and might be responsible for elevated transcription of nearby genes

Using chromatin conformation capture techniques [59] such as HiC [60], it is now well established that the linear genomic DNA is folded into largely conserved topologically associating domains (TADs) in the nuclei of cells [61–63], where distal enhancers interact with promoters via looping over long distances for transcriptional regulation [64, 65]. Therefore, we hypothesize that active exonic TFBSs in a CRM, but not inactive exonic TFBSs in a CRM, must be in close physical proximity to the promoter of the target gene in a TAD, although the promoter may be linearly far away from the exonic TFBSs. To test this, we used a ChIA-PET (chromatin interaction analysis by paired-end tagging) dataset generated in the K562 cell line using an antibody against hypomethylated Pol II at Ser2 [66]. ChIA-PET is a variant of HiC, which combines ChIP-seq with HiC to identify DNA loci that are physically close to sequences bound by the antibody-targeted protein [66]. Pol II hypomethylated at Ser2 stalls at promoters in the preinitiation complex [67], thus the resulting ChIA-PET data are enriched for paired-end reads with one end read mapped to an active promoter and the other end read mapped to an active CRM in close physical proximity to the promoter. As an exonic TFBS tends to be close to the promoter of its host gene, we exclude the promoters of the host genes of exonic TFBSs from analysis (Materials and Methods).

However, since there is still no sufficient knowledge about both active and inactive TFBSs or CRMs even in K562 cells, though being one of the most well-studied ENCODE cell lines, we predicted active and inactive TFBSs in the cells using 793 TF ChIP-seq datasets generated in the cell line (Additional File 2, Table S1). More specifically, we predict a TFBS in a CRM to be active in a cell/tissue type if it overlaps at least a binding peak of a ChIP-ed TF in the cell/tissue type, or, to be inactive, otherwise (Materials and Methods). Obviously, on one hand, the number of predicted active exonic TFBSs in a cell/tissue type depends on the availability of TF ChIP-seq datasets in the cell/tissue type; on the other hand, some of predicted inactive exonic TFBSs might be actually active, but we might lack ChIP-seq data in the cell/tissue type to predict them to be active. Nonetheless, with a relatively large number (793) of TF ChIP-seq datasets available to us (Additional File 2, Table S1), we predicted 328,605 (31.38%) of the 1,047,183 coding TFBSs, 94,428 (72.90%) of the 129,518 5’-UTR TFBSs and 180,193 (24.58%) of the 733,089 3’-UTR TFBSs to be active, and the remaining 68.62% of coding TFBSs, 27.10% of 5’-UTR TFBSs and 75.42% of 3’-UTR TFBSs to be inactive in the cells (Materials and Methods). As shown in Figure 6a, predicted active coding TFBSs (18.44%), 5’-UTR TFBSs (23.78%) and 5’-UTR TFBSs (15.72%) have a significantly higher proportion in close physical proximity to distal promoters than predicted inactive coding TFBS (3.17%), 5’-UTR TFBSs (1.85%) and 3’-UTR TFBSs (2.92%) (p < 2.23x10^-302^ for all comparisons with χ^2^ test), respectively.

**Figure 6.**
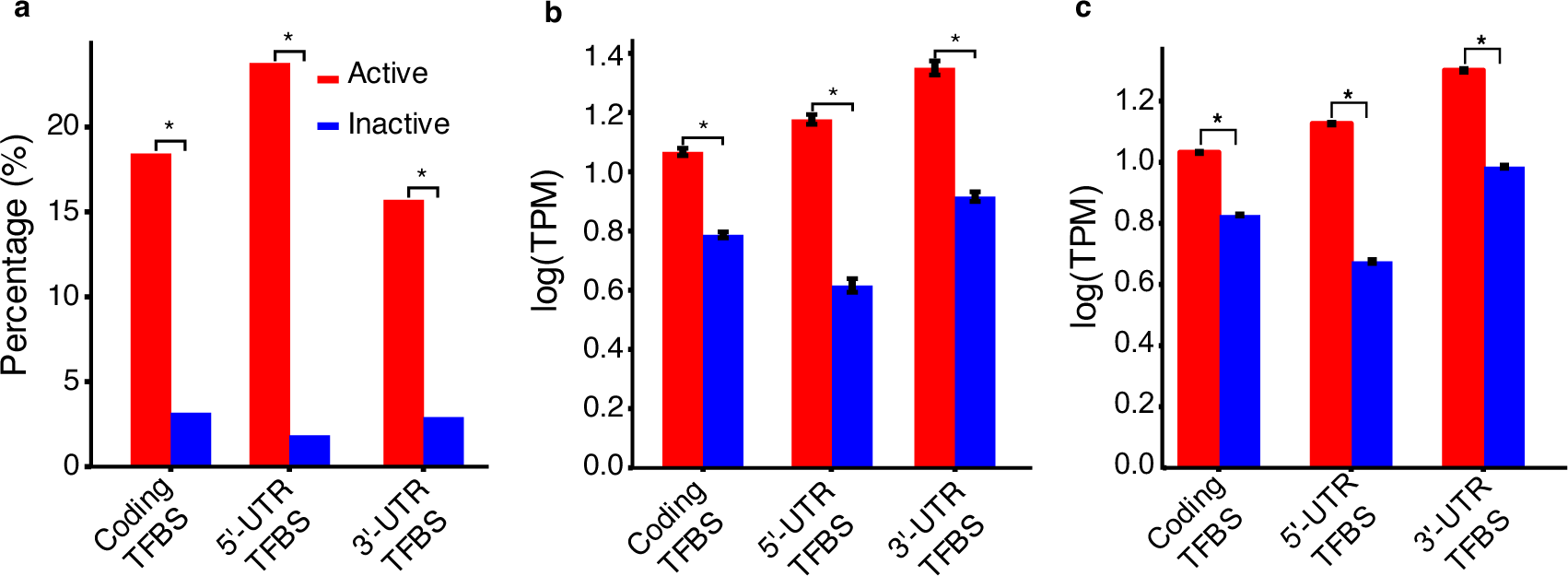
Active exonic TFBSs might enhance the transcription levels of nearby genes. **a.** Percentages of active exonic TFBSs and inactive exonic TFBSs that are in close proximity to non-own promoters in K562 cells. **b.** Transcription levels in K562 cells of genes of promotors that are in close proximity to putative active exonic TFBSs in comparison with those of genes closest to putative inactive exonic TFBSs. **c.** Average transcription levels in the 19 cell/tissue types of genes closest to putative active exonic TFBSs in comparison with those of genes closest to putative inactive exonic TFBSs. Data are shown in mean±SE. The means in (**b**) are for relevant genes in K562 cells, and in (**c**) are for relevant genes in the 19 cell/tissue types. *P<2.23x10^-302^ for all the comparisons was computed using the χ^2^ test (**a**) or Mann Whitney U test (**b, c**).

To see whether these active exonic TFBSs in close proximity to promoters actually regulate the promoters’ genes, we compared transcription levels in K562 cells of genes immediately downstream of promoters that are in close proximity to the putative active exonic TFBSs, with those of genes closest to the putative inactive exonic TFBSs. To avoid confounding factors in the analysis, we excluded exonic TFBSs’ host gene as the closest gene, although an exonic TFBS might regulate its host gene as demonstrated earlier [9, 11, 16, 17]. As shown in Figure 6b, genes immediately downstream of promoters that are in close proximity to putative active coding TFBSs, 5’-UTR TFBSs and 5’-UTR TFBSs have significantly higher expression levels than those closest to putative inactive coding TFBSs, 5’-UTR TFBSs and 5’-UTR TFBSs (p<2.2x10^-302^ for all comparisons with Mann Whitney U test), respectively.

To further verify these observations, we resorted to 19 ENCODE human cell/tissue types in which various numbers of TF ChIP-seq data (Additional File 2, Table S1) as well as RNA-seq data (Additional File 2, Table S2) are available. Similar to the case of K562 cells, in these 19 cell/tissue types, we predicted an average of 31.10%, 78.47% and 28.17% of the 1,047,183 coding TFBSs, 129,518 5’-UTR TFBSs and 733,089 3’-UTR TFBSs to be active, and the remaining 68.90%, 21.53% and 71.83% to be inactive, respectively. As shown in Figure 6c, on average, genes closest to predicted active coding TFBSs, 5’-UTR TFBSs or 3’-UTR TFBSs have significantly higher transcription levels than those closest to predicted inactive exonic TFBSs in the cell/tissue types (p<2.2x10^-302^ for all comparisons with Mann Whitney U test). Taken together, these results strongly suggest that our predicted active exonic TFBSs not only tend to be in close physical proximity to distal promoters, but also might be involved in up-regulation of the nearby genes, and thus might be constituent TFBSs of exonic enhancers.

## Discussion

It has long been known that in addition to encoding amino acids and UTR function-related signals, exons can also encode other information, including splicing enhancers [52–54], overlapping ORFs [68], lincRNA [69] and transcriptional enhancers [9–25]. It has been estimated that up to 25% of codons in the human genome may code for such overlapping functions based on evolutionary constraints on the degenerate positions in synonymous codons [47]. Although there are numerous experimentally verified cases for such dual-uses of codons and UTRs, it is under hot debate as to how prevalent they are in the human genome [30, 70], what evolutionary constraints these codons and UTRs have been subject to in the course of evolution [30, 52, 54, 71], and how it is possible for a sequence to evolve for such two unrelated functions. For instance, Stergachis and colleagues [30] showed that up to 15% codons in 86.9% genes were duons that were predicted using DHSs in 81 human cell/tissue types, and that these duons were under strong negative selection. However, others [33, 34] concluded that duons were largely selectively neutral and thus might not be functional. The discrepancies may result from different methods employed [30, 33, 52] and/or different datasets used [30, 34, 52, 70]. Clearly, the lack of a large experimentally verified positive and negative datasets for exonic CRMs has largely hampered efforts to clarify these contradictory results and address related issues of the dual-uses of exons. Our recent prediction of large sets of CRMCs and non-CRMCs with high accuracy in the human genome provides us an opportunity to address these issues.

Our predicted TFBSs consist of about 40% of the length of CRMC [35], but they might play more critical roles in transcriptional regulation than spacer sequences between them [4]. Moreover, exons only make up less than 10% of the lengths of most exonic CRMs (Figure 1a). Therefore, in this study, we used predicted exonic TFBSs located in experimentally verified exons to address the aforementioned questions for the dual-use of exons. Surprisingly, we found that coding TFBSs, 5’-UTR TFBSs and 3’-UTR TFBSs comprised more than a third of the total length of annotated CDSs, 5’-UTRs, and 3’-UTRs, respectively (Figure 1e), and almost all transcriptional validated protein genes (96%) have at least an exon partially overlapping a CRM (Figure 1c). Therefore, exonic TFBSs and exonic CRMs might be more prevalent than originally thought [30, 47]. Our results suggest that exonic TFBSs as parts of exonic CRMs might be involved in transcriptional regulation of nearby genes. First, we found that predicted active exonic TFBSs are more likely to interact with distal promoters than are predicted inactive exonic TFBSs (Figure 6a). Second, we showed that genes downstream of promoters in close proximity to putative active exonic TFBSs (Figure 6b) as well as genes closest to putative active exonic TFBSs (Figure 6c), had elevated transcription levels compared to those closest to putative inactive exonic TFBSs (Figure 6c).

Evolutionary behaviors of exonic TFBSs strongly support their TF binding functions, in addition to their amino acid coding functions. First, we found that both A/T and C/G at the non-degenerate first (Figures 3a, 3c) and second (Figures 3b, 3d) positions in coding TFBSs were more likely to be under purifying selection than those in non-CRMC CDSs, strongly suggesting that the non-degenerate positions in coding TFBSs might be in dual-use for encoding amino acids in proteins and TF binding interfaces in DNA. Second, degenerate third codon positions in coding TFBSs are more likely under evolutionary constraints (either positive selection for A/T or negative selection for CG) than those in non-CRMC CDSs (Figures 3e, 3f), strongly suggesting that they might be involved in TF binding. Interestingly, both A/T and C/G at degenerate positions in coding TFBSs also are more likely under either negative or positive selections than those in non-exonic TFBSs (Figures 3e, 3f), suggesting that these degenerate positions might also be in dual-use for TF binding and other unknown non-coding functions. This conclusion therefore is in stark contrast to the two earlier contradictory ones drawn by Stergachis et al. [30] and Xing/He [33] using the duons predicted based on DHSs. We [35] and others [36–38, 72] have shown that CRMs predicted based solely on DHSs have high FDRs. Thus, it is highly likely that a high FDR of the earlier predicted duons might lead to the earlier two contradictory conclusions due to the failure to separate A/T and C/G positions for evolutionary analysis by Stergachis et al. [30]. Finally, for the first, to our best knowledge, we found that both 5’-UTR TFBSs and 3’-UTR TFBSs are more likely to be under negative selection than their non-CRMC counterparts (Figures 4a∼4d), strongly suggesting that they might be involved in TF binding.

Interestingly, both 5’-UTR TFBSs (Figure 4b) and 3’-UTR TFBSs (Figure 4d) also are more likely to be under negative selection than non-exonic TFBSs, suggesting that the UTR TFBSs might be in dual-use for TF binding and other UTR-related functions.

The paradox concerning the dual-use of exons is how a DNA sequence could evolve two unrelated functions such as encoding TF binding interface in DNA and amino acids in protein or UTR-related functions. For instance, the amino acid coding function would require the DNA sequence to specify a specific peptide that plays a role in the protein’s function, while the TF binding function would demand the sequence to adopt a specific interface to which the cognate TF could bind. Although an earlier study [12] suggests that nature might solve this paradox by opting exonic remnants from whole-genome duplication for exonic enhancers in some cases, this mechanism cannot explain the prevalent dual-use of exons as we show in this study and others reported earlier [47]. However, our findings might provide a more general answer to the puzzle. First, at least a quarter of the non-degenerate first and second codon positions in non-CRMC CDSs are selectively neutral or nearly so (Figures 3c, 3d), thus they could potentially evolve into coding TFBSs without detrimental effects on protein functions. Interestingly, the proportion of selectively neutral non-degenerate positions in coding TFBSs is much smaller than in non-CRMC CDSs (Figures 3c, 3d), suggesting that once originally neutral non-degenerate positions become coding TFBSs, they are under purifying selection. This might explain why the proportions of selective neutrality in non-degenerate positions in coding TFBSs are lower than those in non-CRMC CDSs (Figures 3c, 3d). Second, amino acids encoded by coding TFBSs tend to be located in structurally and functional less critical loops, and avoid structurally and functionally more important helices and strands (Figures 5a and 5b), thereby reducing detrimental effects of evolving codons into coding TFBSs. Third, the preferential locations of coding TFBSs at 5’-ends and 3’-ends of a concatenated CDS of a gene (Figure 1f) suggest that coding TFBSs tend to encode amino acids at N-terminus and C-terminus of a protein. These termini probably have less crucial functions than those in the middle of the peptide. Fourth, 5’-UTR TFBSs tend to be located at the middle of 5’-UTRs (Figure 1f), and avoid the two ends where 5’-UTR function-related sequences are encoded, such as transcription start sites (TSSs), ribosome entry sites and upstream open-reading frames [55, 73]. Finally, 3’-UTR TFBSs tend to appear at the two ends of 3’-UTRs (Figure 1f), and avoid the middle where 3’-UTR function-related sequences might be coded, such as polyadenylation sites, miRNA response elements and RBP (RNA binding protein) binding sites [56]. To summarize, all these results, strongly suggest that a high level of prevalent dual-uses of both CDS and UTRs are possible.

The next interesting question is, when only 4.1% of the human genome code for exons [74] and the remaining 95.9% are NESs that can be potentially used to encode CRMs, why are the sparse exons exploited to encode TFBSs in CRMs? Our finding that active coding TFBSs, 5’-UTR TFBSs and 3’-UTR TFBSs tend to be in close physical proximity to distal promoters in the 3D space of the nuclei may provide an explanation. When chromatins are folded into relatively conserved 3D structures [61–63], linearly distal enhancers and target promoters are brought in close physical proximity in compartments such as TADs that typically span around 1 Mbp [64, 65, 75]. Since functionally related genes tend to cluster together in linear DNA, while gene deserts may harbor no genes [62, 76], it is highly likely that there is not enough NESs in close proximity to a promoter to function as tis enhancer in a TAD [77]. In such a scenario, a few nucleotides in a CDS or UTR in close proximity to the promoter may well likely evolve into coding TFBSs, if they code for less important amino acids such as those in some loops, termini of proteins, or a less critical part of UTRs. In this regard, dual-uses of some exons might be unavoidable, because a NES is physically unavailable to adopt the CRM role although it might be more evolvable. Therefore, it appears that nature chooses less critical exons for TFBSs, thereby avoiding the dilemma of evolving a sequence for two unrelated functions.

Based on our findings in this study, we propose a parsimonious model for how a stretch of codons in an exon could evolve into a coding TFBS. As illustrated in Figure 7a, when no NES is in close proximity to a (potential) promoter (e.g., P’) in a TAD due to its limited space to hold more NESs, nature might choose a stretch of less critical codons in an exon (e.g., E4) that is physically close to the (potential) promoter to evolve into a desired TFBSs. Such possibility would increase if the non-degenerate positions of this stretch of codons (e.g., codons 1∼4 in Figure 7b) match nucleotides in some positions in the desired TFBS, because such adoption of a TFBS role would not alter the encoded amino acids. This model predicts that these non-degenerate positions would be subject to double negative selections, and thus, are more likely to be under negative selection than those in non-CRMC CDSs. This is exactly what we observe as shown in Figures 3c and 3d. At the same time, nature would choose the degenerate positions of the codons to either mutate to the desired nucleotides (codons 1 and 2 in Figure 7b) or remain the same if they match the desired nucleotides in the positions of the desired TFBS (codon 3 in Figure 7b). This model predicts that, in the former case, the mutated degenerate positions would be under positive selection, while in the latter case, the matched positions would be subject to negative selection. Since C/G have high contents at the degenerate third codon positions (Figures 2a, 2b) [49], the chance for C/G at these positions to match C/G at the positions in the desired TFBSs could be high, and thus, no mutation is needed. On the other hand, as A/T have low contents at the degenerate third codons positions (Figures 2a, 2b) [49], the chance for A/T at these positions to match A/T at the positions in the desired TFBSs could be low, and more likely, C/G to A/T substitutions are needed to evolve into the desired TFBS. Therefore, our model predicts that C/G and A/T at the degenerate third codon position are more likely to be under negative and positive selections, respectively, than those in non-CRMC CDSs. This is exactly what we see as shown in Figure 3f. Therefore, our model nicely explains the observed evolutionary behaviors of both A/T and C/G at the non-degenerate and degenerate positions in coding TFBSs compared with those in non-CRMC CDSs. Similarly, parsimonious models of how a stretch of less critical nucleotides in a UTR that is in close physical proximity to a (potential) promoter would evolve into a UTR TFBS can be envisaged based on our findings presented in this study.

**Figure 7.**
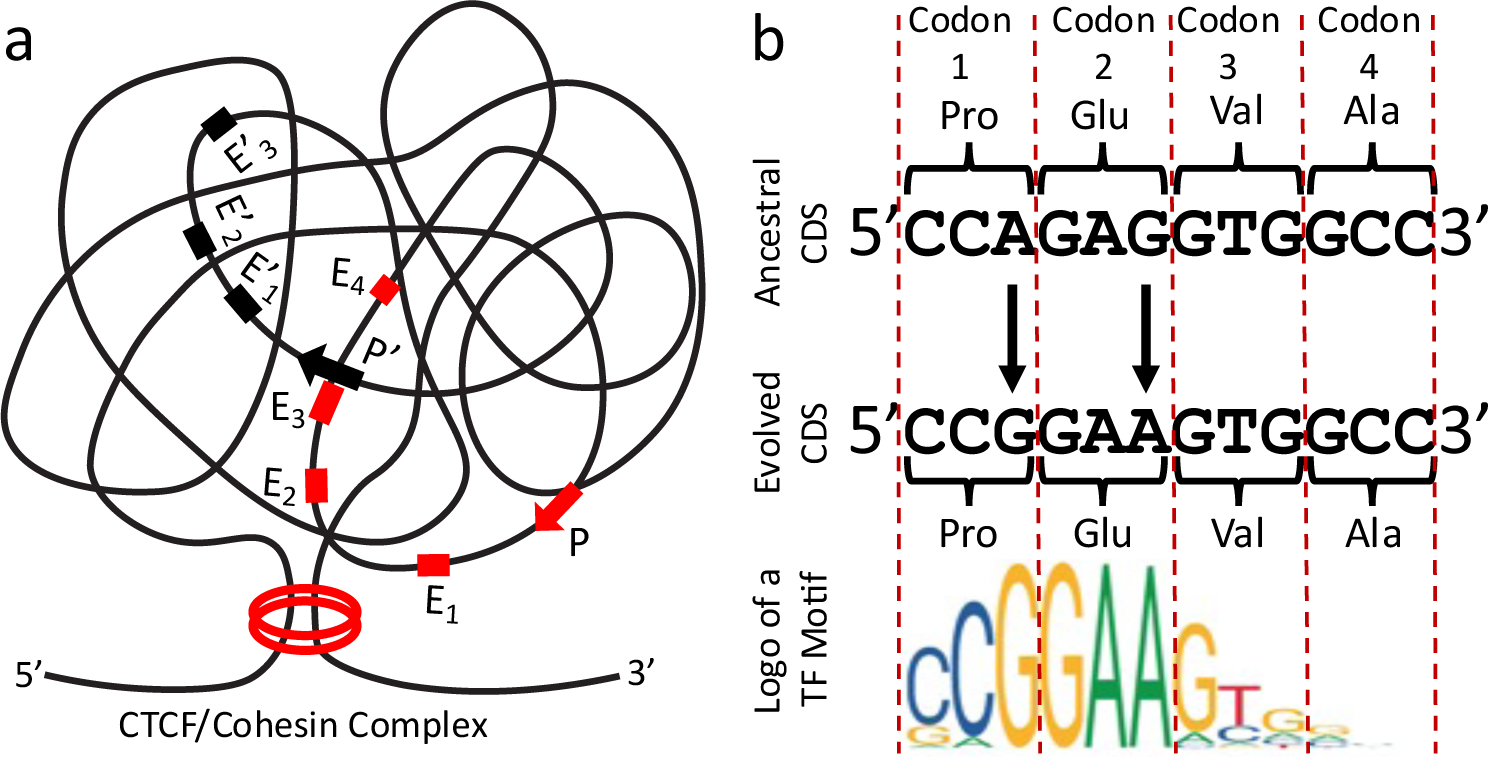
A parsimonious model for how a stretch of codons evolve into a TFBS. **a.** A carton of a TAD locked by a CTCF/cohesin complex. Displayed are two hypothetical genes under the control of promoters P and P’, containing exons E_1_∼E_4_ and E’_1_∼E’_3_, respectively. Exon E_3_ is physically close to promoter P’, while no NES is physically close to P’. If a stretch of codons in E_3_ encode a less critical short amino acid sequence in the protein, and/or if some non-degenerate first and second codon positions match a desired TFBS, then this stretch of codons could evolve into a desired TFBS without little or no detrimental effects on the protein functions. **b.** A hypothetical parsimonious scenario in which a stretch of four codons in an ancestral CDS evolved into a TFBS of a TF recognizing a motif with the displayed logo through two synonymous substitutions at the degenerate positions of codons 1 and 2, without changing encoded amino acid sequence. Since the nondegenerate positions of codons 1∼4 as well as the nondegenerate position of codon 3 match the desired TFBS, the model predicts that they would be subject to negative selections. As the non-degenerate positions of codons 1 and 2 are mutated to the desired nucleotides, the model predicts that they would be subject to positive selections.

## Conclusions

Exonic TFBSs occupied at least a third of the total exon lengths, and 96.7% of genes had exonic TFBSs, suggesting that they might be more prevalent than originally thought. Both A/T and C/G in exonic TFBSs are more likely under evolutionary constraints than those in non-CRMC exons, indicating that they are likely in dual-use. Exonic TFBSs in codons tend to encode loops rather than more critical helices and strands in protein structures, while exonic TFBSs in untranslated regions (UTRs) tend to avoid positions where known UTR-related functions are located. Moreover, active exonic TFBSs tend to be in close physical proximity to distal promoters whose immediately downstream genes have elevated transcription levels, suggesting that they might be involved in transcriptional regulation of target genes. It is highly possible that less critical positions in an exon that is physically close to a promoter can evolve into a TFBS when no non-exonic sequences are physically available to the promoter.

## Materials and Methods

### Datasets

We downloaded predicted CRMCs and non-CRMCs from the PCRMS database (https://cci-bioinfo.uncc.edu/) [78], and extracted putative CRMs with a p-value < 5x10^-6^ to minimize FDR, resulting in 428,628 CRMs containing 38,507,543 potentially overlapping putative TFBSs. The human genome assembly version GRCh38.p13 was used as the reference genome. Annotations (CDSs, 5-UTRs and 3’-UTRs) of genes and verified transcripts were downloaded from Ensembl Release 100 (https://useast.ensembl.org/index.html). For each gene, the longest annotated transcript (agreed by different databases, with support from mRNA data) was selected for CDS assignments. Protein structures were download from the Protein Data Bank (PDB) [79]. The chromatin interaction analysis by paired-end tagging (ChIA-PET) dataset of the K562 cell line [66] was downloaded from the GEO database [80] with the access number GSE33664. TF ChIP-seq datasets in 19 human cell/tissue types (Additional File 2,Table S1) were downloaded from the Cistrome database [81]. RNA-seq datasets in the 19 human cell/tissue types (Additional File 2, Table S2) were downloaded from the ENCODE data portal (https://www.encodeproject.org/about/data-access/).

### Definition of degenerate positions and non-degenerate positions of the codons

We consider genome nucleotide positions at the third codon positions of the 21 synonymous codon sets to be degenerate, including 12 sets of two-fold degenerate positions, i.e., Arg(AG[AG]), Asn(AA[CT]), Asp(GA[CT]), Cys(TG[CT])), Gln(CA[AG]), Glu(GA[AG]), His(CA[CT]), Leu(TT[AG]), Lys(AA[AG]), Phe(TT[CT]), Ser(AG[CT]) and Tyr(TA[CT]); one set of three-fold degenerate positions, Ile(AT[ACT]), and eight sets of four-fold degenerate positions, i.e., Ala(GC*), Arg(CG*), Gly(GG*), Leu(CT*), Pro(CC*), Ser(TC*), Val(GT*), Thr(AC*). We consider genome nucleotide positions at the first codon positions of all the sense codons to be non-degenerate, except for Arg ([AC]GA), Arg ([AC]GG), Leu ([CT]TA) and Leu ([CT]TG), because they are two-fold degeneracy. We also consider genome nucleotide positions at the second codon positions of all sense codons to be non-degenerate.

### Assignment of secondary structure types

For proteins with known structures in PDB, we assigned secondary structure types to amino acid sequences using the DSSP program [82] following the widely used convention: H (α-helix), G (310-helix) and I (π-helix) states as helix type; E (extended strand) and B (residue in isolated β-bridge) states as strand type and all the other states as loop type. For proteins without known structures, we assigned secondary structure types as follows. We first generated a non-redundant protein sequence set (less than 30% sequence identity) for all annotated proteins in the human genome using CD-HIT [83]. If a protein in the dataset has a highly homologous protein with known structure in PDB with at least 50% coverage and 80% sequence identity, the secondary structure types of the template protein were copied to the target protein. If no homologous structures were found in PDB, RaptorX, a highly accurate secondary structure prediction program, was applied to predict secondary structure types with default settings [58].

### Assignment of conservation scores

The phyloP [32] scores of each nucleotide position in the human genome were downloaded from the UCSC Genome Browser database [84].

### Prediction of active and non-active TFBSs in cell lines

If a TFBS entirely falls in a 1,000bp region centering on the summit of any TF ChIP-seq binding peak in a cell line, we predict the TFBS to be active in the cell line. Otherwise, we predict the TFBS to be inactive in the cell line. Clearly, the completeness of prediction of active TFBSs in a cell/tissue type depends on the number of available ChIP-seq datasets for active TFs in the cell/tissue type.

### Validation of exonic TFBSs using chromatin interactions

We calculated significant interactions between two loci using ChiaSig [85] based on the ChIA-PET paired-end reads from K562 cells [66]. We then counted the number of predicted active and inactive exonic TFBSs overlapping loci in close proximity to the promoter that was not the immediate upstream to host gene of the exonic TFBSs, to exclude the promoters of the host gene of exonic TFBSs.

### Statistical analysis

Kolmogorov-Smirnov(K-S) test, Mann Whitney U test or χ□ test was used to evaluate statistically significant levels of hypothesis tests as indicated in the text and figure legends.

## Supplementary data

Additional File 1: Figure S1, examples of experimentally validated exonic enhancers. Additional File 2: Table S1, summary of the TF ChIP-seq datasets in the 19 human cell/tissue types; Table S2, summary of the RNA-seq datasets in the 19 human cell/tissue types.

## Authors’ contributions

J.C. performed the data analysis. P.N. and M.N performed CRM and TFBS predictions. Z.S and J.T.G conceived of and supervised the study. Z.S. wrote the manuscript. All author(s) read and approved the final manuscript.

## Availability of data and materials

All datasets and predicted results as well as Python and R scripts for data processing and analyses are available upon request.

## Declarations

### Ethics approval and consent to participate

Not applicable.

## Acknowledgements

We would like to thank Dr. Way Sung for stimulating discussions and comments on the manuscript.

