## Supplementary material for "Prevalent uses and evolution of exonic regulatory sequences in the human genome": Figure S1

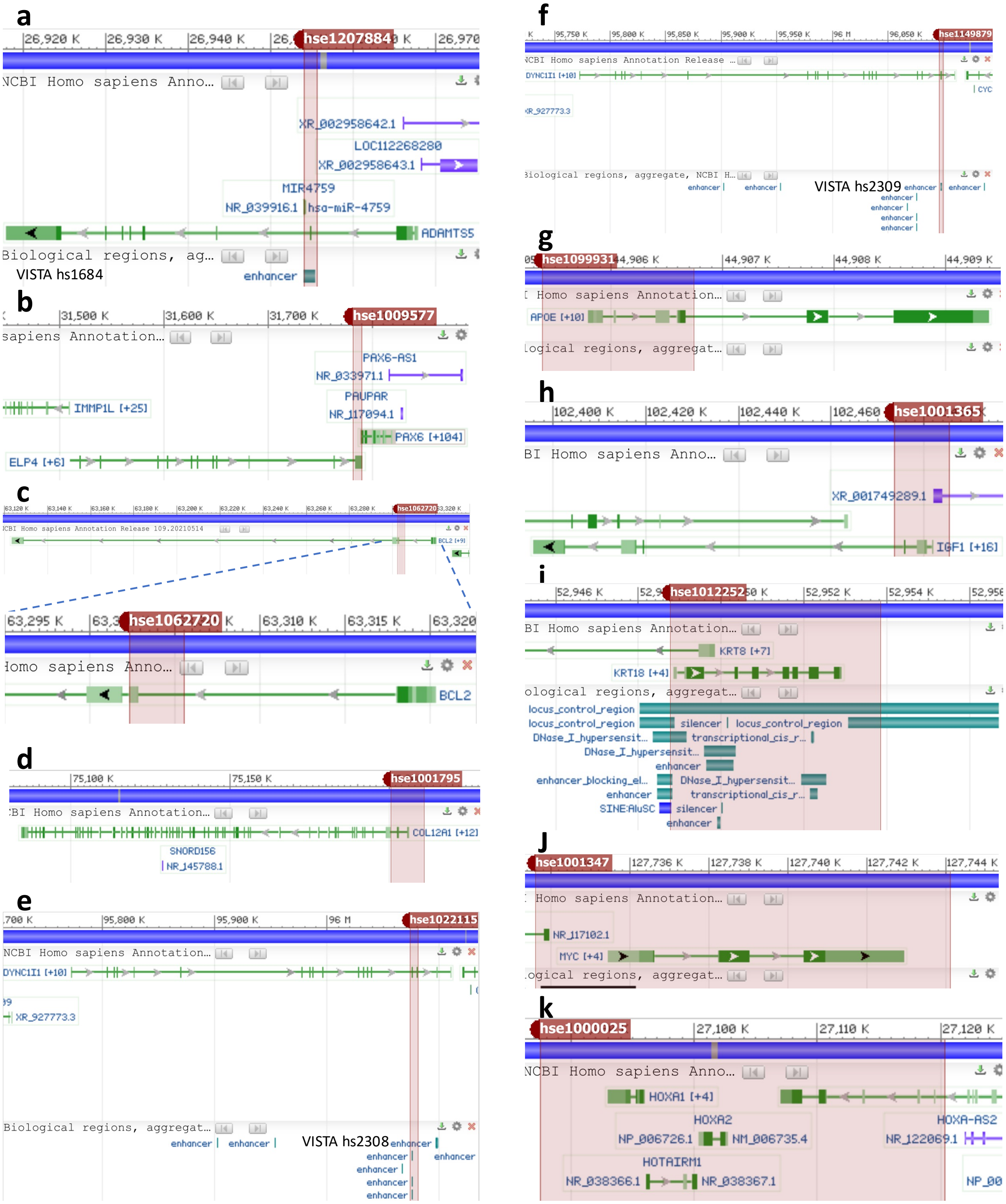


**Figure S1.** NCBI genome viewers of examples of the predicted CRMs overlapping experimentally characterized exonic enhancers. **a.** Predicted CRM hse1207884 (1,486bp) spans an experimentally validated exonic enhancer containing the second exon of the *ADAMTS5* gene that regulates its own transcription[11]. Hse1207884 also overlaps VISTA enhancer hs1684 [43]. **b.** Predicted CRM hse1011628 (7,096bp) is a supper-enhancer that spans an experimentally validated exonic enhancer (not shown) containing the last exon of the *ELP4* gene that regulates the transcription of the downstream gene *PAX6* [12, 86]. **c.** Predicted CRM hse1062720(3,111bp) spans an experimentally validated exonic enhancer (not shown) containing the second exon of the *BCL2* gene that regulates its own transcription [16]. **d.** Predicted CRM hse1001795 (10,032bp) is a supper-enhancer that spans an experimentally validated exonic enhancer (not shown) containing the 1st and 2nd exons of the collagen type XII alpha 1 chain gene *COL12A1*[24]. **e.** Predicted CRM hse1022115 (6,202bp) is a supper-enhancer that spans an experimentally validated exonic enhancer containing the 15th and 16th exons of the *DYNC1I1* gene that interacts with and regulates the promoter of the *DLX5/6* gene[13]. Hse1207884 also overlaps VISTA enhancer hs2308 [43]. **f.** Predicted CRM hse11149879 (2,816bp) spans an experimentally validated exonic enhancer containing the 17th exon of the *DYNC1I1* gene that might regulate the *DLX5/6* gene[27, 28]. Hse1149879 also overlaps VISTA enhancer hs2309 [43]. **g.** Predicted CRM hse1099931(1,353bp) spans an experimentally validated exonic enhancer (not shown) containing the 3rd and 4th exons of the *APOE* gene[25]. **h.** Predicted CRM hse1001365(11,368bp) is a super-enhancer that spans an experimentally validated exonic enhancer (not shown) containing the 1st exon of the *IGF1* gene[23]. **i.** Predicted CRM hse1012252 (5,036bp) is a super-enhancer that spans the entire *KRT18* gene locus. hse1012252 overlaps a 10-kb locus control region (LCR)[87], an upstream LCR and an downstream LCR [88]. hse1012252 also overlaps an experimentally validated exonic enhancer containing the sixth exon that interacts with the enhancer in the first intron to regulate KRT18 transcription [9]. **j.** Predicted CRM hse12001347(10,483bp) is a supper-enhancer that spans the entire *MYC* gene locus. hse12001347 overlaps an experimentally validated exonic enhancer (not shown) containing the 1st exon of the *MYC* gene that regulates its own transcription [18]. **k.** Predicted CRM hse1000025 (32,725bp) is a super-enhancer that runs through multiple *HOX* genes and known enhancers regulating expression patterns of the HOX gene cluster[89-91], including the exonic enhancer (not shown) that spans the two exons of the *HOXA2* gene[10, 17].
